# Origin, exposure routes and xenobiotics impart nanoplastics with toxicity on freshwater bivalves

**DOI:** 10.1101/2022.10.27.514081

**Authors:** Adeline Arini, Sandra Muller, Véronique Coma, Etienne Grau, Olivier Sandre, Magalie Baudrimont

## Abstract

Various environmental aged plastic wastes were collected in the environment and crushed to the nanometric scale to get a mix of nanoplastics (NPs) of different natures – mostly polyolefins (PE, PP), polyesters (PET) and polyvinylics (PS and PVC) – and undefined shapes (noted NP-L, mean hydrodynamic diameter at 285 nm). We aimed to test the toxicity of NPs of environmentally relevance on freshwater bivalves and compare results to commonly used styrenic NP-PS (206 nm). *Corbicula fluminea* were exposed to four different conditions with NPs (0.008 to 10 μg/L), for 21 days and kept under depuration conditions for 21 additional days: 1) waterborne exposure to NP-L, 2) diet borne exposure to NP-L, 3) synergic waterborne exposure to NP-L and AlCl_3_ salt (1 mg/L), 4) waterborne exposure to NP-PS. Enzyme activities, gene expressions and behavioural tests were assessed. Trophic and synergic exposures with Al triggered more gene modulations than direct exposure to NP-L (namely on *12s, atg12, gal, segpx, p53* and *ache*). NP-PS were also more harmful than NP-L, but only at high concentrations (10 μg/L). The effects of each treatment lasted until 7 days of depuration and no more gene inductions were observed after 21 days in clean water. Very few effects were shown on phenol-oxidase (PO), and glutathione S-transferase (GST). However, the inhibition of acetylcholinesterase (AchE) was concomitant with an increase of the filtration activity of bivalves exposed to NP-L (trophic route) and NP-PS, suggesting neurotoxic effects. By disturbing the ventilatory activity, NPs could have direct effects on xenobiotic accumulation and excretion capacities. The results point out how the structure, aging, exposure routes and additional xenobiotics can interact with adverse outcomes of NPs in bivalves. These findings underline the importance to consider naturally aged environmental NPs in ecotoxicological studies rather than synthetic latexes, *i*.*e*. crosslinked nanospheres prepared from virgin polymers.

This manuscript presents the first data of toxicity on freshwater organisms exposed to nanoplastics coming from natural sources. Whereas the majority of papers are dealing with non-environmentally representative plastics (mainly commercially-available polystyrene latexes) to evaluate nanoplastic effects on organisms, this study develops methods to prepare model nanoplastics from plastic wastes collected in rivers, and to assess their real adverse effects on aquatic organisms. Our results show significant differences between the inflammatory effects triggered by nanoplastics coming from natural sources and polystyrene nanobeads. This work suggests that the data published so far in the literature may underestimate the toxicity of nanoplastics spread into the environment on the aquatic organisms at the bottom of the food chain, which might consequently impart halieutic resources on the long term.

## INTRODUCTION

The increasing production of plastics has resulted in millions of tons of plastic wastes spread into the environment. Large plastic debris can break down into micro- and nanometric particles. Several definitions exist to define nanoparticles of plastics, commonly called “nanoplastics” (NP). They correspond to plastic particles whose size is comprised between 1 and 1000 nm, resulting mainly from the degradation of larger plastic particles and having a colloidal behaviour in the environment ^1-3^. In addition to the degradation of plastic debris, another important source of NPs arises from textile fibres rejected by washing machines and the inability of waste water treatment plants to efficiently filter all these particles before releasing water into the aquatic environment ^4^.

Characterizing these particles at the nanometric scale and at very low concentrations in the environment requires solving several analytical issues. One of the biggest challenges relies on the lack of methods for the quantification of NPs, most of which are at their infancy, making almost impossible to obtain real concentrations of NPs in the environment. It is thus difficult until now to know the sources of emission, the fate and responsiveness of NPs. Therefore ecotoxicology studies carrying out on NP toxicity coming from the environment are still very scarce.

NPs’ nanometric properties greatly differentiate them from the rest of plastic wastes ^5^. The hydrophobicity of NPs added to their important surface/volume ratio facilitates their interactions with system components such as organic matter, micro-organisms, dissolved metals and organic pollutants ^6^ as well as the adsorption of hydrophobic organic and inorganic compounds present in the environment ^7-9^. As a consequence, NPs can act as vectors of contamination into the environment and even into aquatic organisms, which ingested or absorbed them, since they can cross biological barriers (cells, tissues, etc.) ^10^.

Several studies have reported their toxic effects, in particular on growth, locomotion, energetic metabolism, associated with the production of reactive oxygen species (ROS) that can lead to death ^11-15^. These important interactions with the aquatic life suggest that NPs can play a key role in the functioning of ecosystems. However, to date more than 95% (source Scopus, publications from 2017-2022) of the studies relie to only one type of commercial NP: polystyrene (PS) under the form of commercial latexes, which are spherical and chemically crosslinked particles, without taking into account the variability of the plastic polymers produced and released into the environment.

Moreover, most of the studies in ecotoxicology conducted to date are performed at unrealistic doses of NP exposures, in the range of the mg/L ^1, 16-19^.

NPs are not only potentially toxic for aquatic life, but they can also act as carriers of chemicals, which may modify their behaviour, their reactivity and induce the release of contaminants farther away from the original sources. Coupling NPs with other types of contaminants is a very innovative approach since no data exists so far about the potential synergistic effects of NPs with associated contaminants in freshwater species. It is also worth noting that recovery phases are rarely included in experimental studies and very scarce data exist about recovery capacities of organisms ^20^.

Our study aimed to use plastic wastes collected in the Leyre river (South-West of France), because it is of great ecologic, touristic, and economic importance. The intense nautical activities in Summer can be sources of various plastic wastes in the Leyre, potentially carrying adsorbed contaminants. Different collected plastic wastes were mixed and crushed to the nanometric scale before being used under laboratory conditions to mimic environmental NPs.

The objectives of this study were to assess the toxicity of these “environmental NP” suspension and compare it with nanoparticles made from virgin polystyrene (NP-PS) commonly used in the literature. We tested different contamination routes (water- and diet borne exposures) on a freshwater bivalve, *Corbicula fluminea*. We used environmentally realistic concentrations (0.008 to 10 μg/L) ^21, 22^ with or without synergetic contamination by aluminium cations (Al^3+^), naturally found in the Leyre river from the leashing of aluminosilicate minerals. Different outcomes were tested, from molecular to individual levels to get an integrative overview of NPs’ toxicity. The exposure was followed by a depuration phase to explore recovery capacities of bivalves after the different exposure treatments.

This study should bring the first outcomes about the toxicity of environmentally realistic NPs on a freshwater species, and help understanding to which extend NPs’ toxicity may depend on the exposure route, and potential interference with associated contaminants. It also investigates for the first time the recovery capacities of bivalves after different NP exposures, at the gene and enzymatic levels.

## METHODS

### Nanoplastic preparation and characterization

Plastic macro-wastes were collected in different sites on the banks of the Leyre river (South-West of France) in July 2020, by the citizen association “La Pagaie Sauvage” (https://lapagaiesauvage.org). After alkaline treatment (KOH 3 M for 48 h at 20°C) to remove any adsorbed biological organic matter, a random mix of plastic wastes was crushed into small pieces in a mill grinder (IKA-A10) filled with liquid nitrogen to make the plastics more brittle. The millimetric powder was then crushed in liquid nitrogen (centrifugal grinder Retsch ZM 200) and sifted using a final 80 μm final sieve. The fine powder was resuspended into milliQ water and sonicated for 10 min with an ultrasound bath. The suspension was filtered with glass-fibre syringe pre-filters (Nalgene, pore size of 1 μm) to obtain a nanoscale suspension called NP-L.

Polystyrene pellets were purchased from Sigma Aldrich (CAS 9003-53-6, M_w_=200 kg/mol, Ð=M_w_/M_n_=2.2 according to size exclusion chromatography, see Figure S1). PS pellets were dissolved in tetrahydrofuran solvent (THF purchased from VWR, CAS 109-99-9) (10 mg PS/mL THF) for 10 min under agitation. Then milliQ water (10:1 volume ratio) was added under agitation to perform the “solvent shift” or “nanoprecipitation” process of PS chains into nanoparticles, and left under stirring for 2 h at 50°C, according to standard procedure as in Schubert et al.^23^. The nanoprecipitated PS suspension was sonicated for 10 min in an ultrasound bath and filtered with 1 μm glass-fiber syringe pre-filters to obtain a nanoscale suspension called NP-PS. The suspension was left open under a hood for a week before use, to enable total evaporation of the THF solvent. Analysis by GC-MS was performed on the filtrate of an aliquot of the NP-PS suspension without detectable residual presence of THF inside the sample (data not shown).

The electrokinetic zeta-potential of particles was measured by phase analysis light scattering zetametry (PALS, Malvern NanoZS) to determine the surface charge of NPs. The hydrodynamic size of the particles was assessed by dynamic light scattering (DLS, Vasco Flex instrument from Cordouan technologies), working at a backscattering angle of 165° and using both the Cumulant fitting method (Table 1) and the Pade-Laplace multimodal analysis (Figure S2). NP suspensions were imaged by transmission electron microscopy (TEM) with samarium acetate staining with a Hitachi H7650 electron microscope operated at 80 kV. The weight concentrations of NPs in the suspensions (dry extracts) were measured by thermogravimetry analysis (TGA, Q50, TA instrument, Figure S4). The polymers present in NP suspensions was analysed by Fourier-transform infrared spectroscopy (FTIR, Bruker Vertex 70, Figure 1).

**Table 1:** Plastic nanoparticle characteristics

|  | Stock solution<br>concentration (mg/L) | Z-average diameter<br>(nm)/PDI | Zeta potential<br>(mV) | Polymer composition |
| --- | --- | --- | --- | --- |
| NP-L | 147 | 285/0.25 | -32 ± 6 | Polypropylene and<br>Polyethylene mostly |
| NP-PS | 420 | 206/0.17 | -44 ± 6 | Polystyrene |

**Figure 1:**
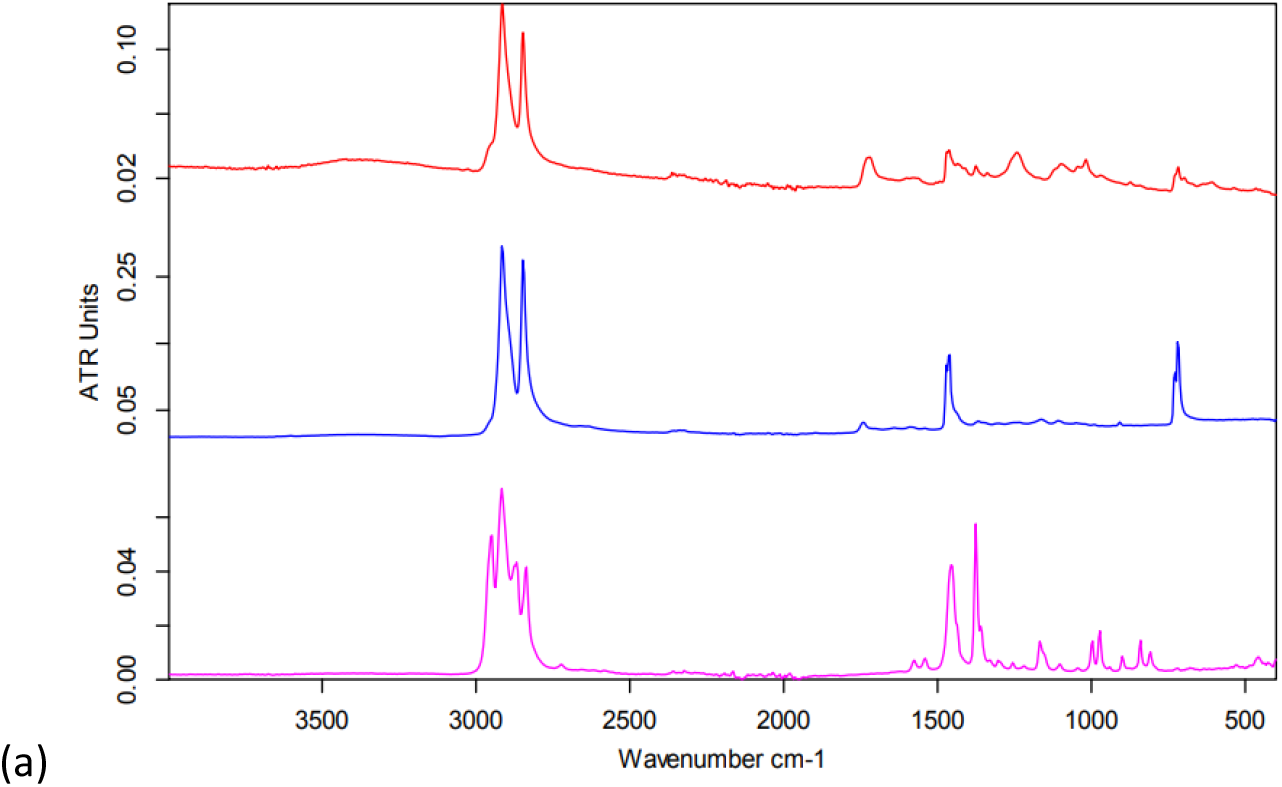
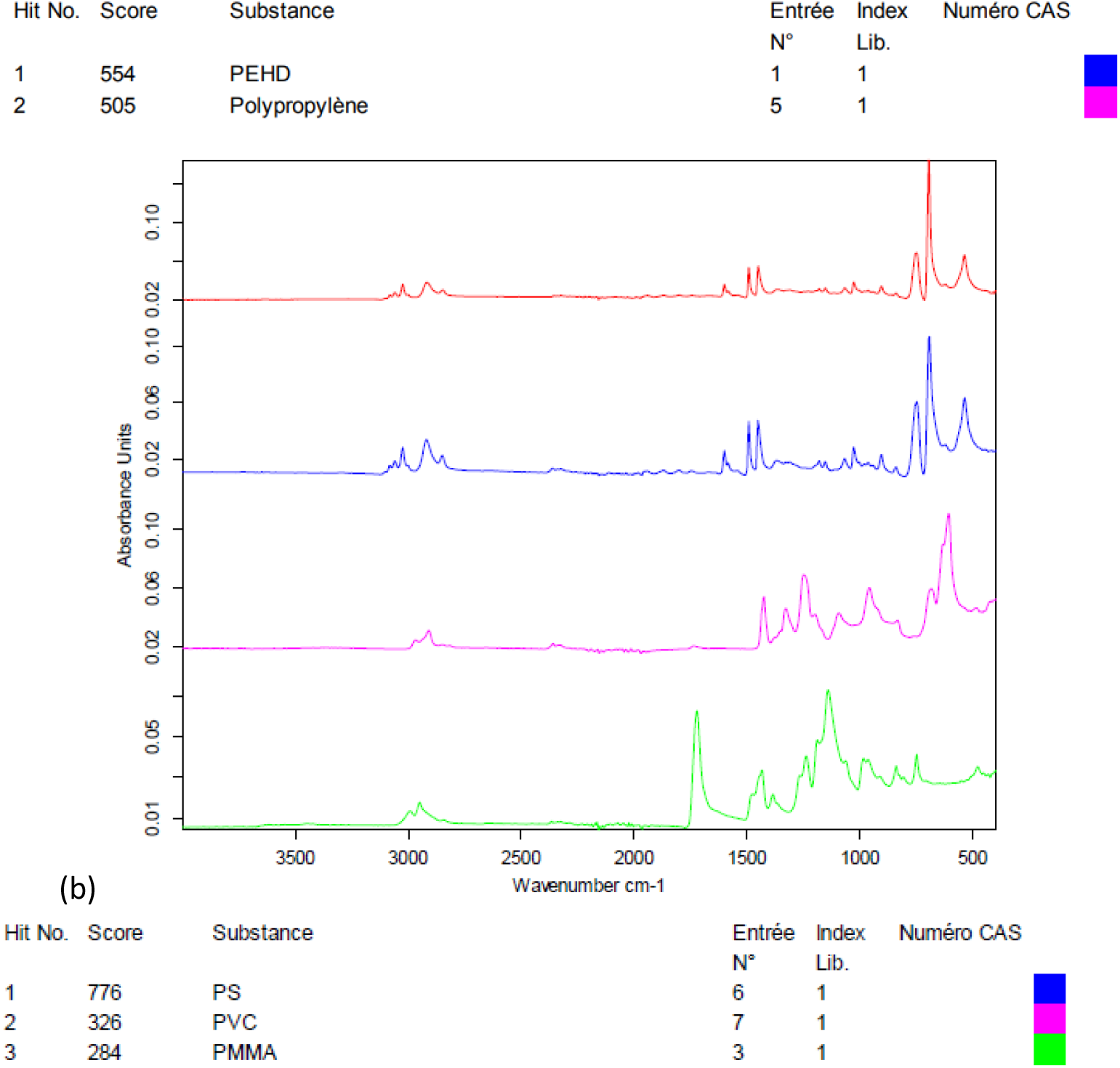
Fourier transform infrared (FT-IR) spectra of the environmental nanoplastic suspension NP-L (in red) best matching the tabulated spectra of PE and PP in the database; b) FT-IR spectra of the model suspension NP-PS (in red) compared to standard vinylic polymers in the database (PS, polyvinyl chloride (PVC) and poly(methyl methacrylate) (PMMA)).

### *Corbicula fluminea* collection and acclimation

Around 300 Asiatic clams *Corbicula fluminea* were collected in a small tributary of the Isle River in Montpon-Ménestérol, considered as clean. Clams were kept in clean water for 2 weeks for acclimation, until used for experiments. Clams were fed with the green algae *Scenedesmus subspicatus* (70 000 cells/individual/L) every other day.

### Experimental design

Glass vessels were filled with 1 L of dechlorinated tap water at 15°C and oxygenated with aeration pumps two days before the start of the experiment. Light was provided by artificial neon lamps (photoperiod 12:12). The day before the start of the experiment, 8 clams were introduced in each experimental unit to get used to the experimental conditions.

The day when the experiment started, contaminants were spiked directly into tanks intended for waterborne exposures. Three concentrations of NP-L and NP-PS were used: 0.008, 1, 10 μg/L. Concentrations were chosen according to published data of microplastic concentrations in deep sea and oceanic gyres ^21, 22^ since no data exists so far for NP concentrations into the environment. Clams were exposed to NP-L with or without an additional contaminant: aluminium salt (AlCl_3_), at 1 mg/L.

Clams dedicated to diet borne exposures were fed with algae solutions that had been contaminated with 0.008, 1 or 10 μg/L of NP-L under agitation, 48 h before the start of exposure. The day of the start of the experiment, contaminated algae were introduced into the experimental units (70 000 cells/individual/L).

Clams dedicated to waterborne exposures were fed with *Scenedesmus subspicatus* (70 000 cells/individual/L) 2h before the exposure started, to avoid contamination by the trophic route once NPs were spiked into the experimental units.

There were four experimental replicates per condition, according to the following denominations:

- Controls
- Waterborne exposure with NPs from the Leyre River: NP-L
- Waterborne exposure with PS: NP-PS
- Synergic waterborne exposure with NPs and Al: Al + NP-L
- Waterborne exposure with Al: Al
- Diet borne exposure, with *Scenedesmus subspicatus* contaminated with NP-L: Diet NP-L

Temperature, Al concentrations and water levels were checked daily to be adjusted, if necessary. In the waterborne conditions, clams were fed every other day, 2 hours before water was changed. Dechlorinated water was renewed every other day in each tank, and contamination was added after water renewal (either spiked contaminants or contaminated algae) to be kept as close as possible to nominal concentrations.

No mortality was observed during the experiment. Clams were sacrificed after 7 (T7) and 21 days (T21) of exposure to NPs and/or Al, and after 7 (D7) and 21 days (D21) of depuration. Gills and visceral mass were dissected. Organs from two individuals were pooled to get enough mass, and split for the different analyses: Al bioaccumulation, enzyme activities and gene expression. Gills and visceral mass dedicated to transcriptomic and enzymatic tests were stored at −80°C before analysis. Bioaccumulation and enzymes activities were performed on samples from T7, T21 and D7 (since no significant differences were shown after 7 days of depuration), whereas transcriptomic analyses were continued until D21.

### Aluminium bioaccumulation

Samples collected for Al bioaccumulation were dried at 40°C for 48h before mineralization (1 mL of nitric acid at 100°C for 3 hours). A volume of 6 mL of ultrapure water was added to digestates and samples were stored at 4°C before analysis. Aluminium was analysed by atomic absorption spectrometry 240Z AA (AAS, Agilent Technologies). The validity of the Al titration method was periodically checked with an intern laboratory certified reference material.

### Enzyme assays

#### Protein assay

Samples were homogenized with a FastPrep®-24 homogenizer (3 times for 40 s), in Tris buffer (pH 7.4) and centrifuged at 12 000 rpm for 15 min to retrieve the supernatant used for assays. Protein concentrations were measured using the Bradford Assay. Briefly, 5 μL of each homogenized samples were added to 250 μL of Coomassie blue reagent. The samples were incubated for 10 minutes at room temperature before the absorbance was read in a microplate spectrometer (BioTek EPOCH) at 620 nm. A standard curve was built using BSA (Bovin serum albumin) dilutions from 0.1 to 1000 μg/mL. The slope of the standard curve was used to calculate the protein concentration of samples. A sample concentration of 300 μg/mL was used for each enzymatic assay (4 analytical replicates per sample).

#### Phenol oxidase assay (PO)

The PO activity was measured as described in Le Bris et al ^24^. A volume of 50 μL of gills supernatant at 300 μg/mL was added to a 96-well microplate. Samples were incubated 10 minutes with 50 μL of Tris-HCl buffer (0.1 M at pH 8). A solution of L-3,4-dihydroxyphenylalanine was prepared (*L*-Dopa, 20 mM in Tris-HCl buffer) which is a common substrate to the three PO subclasses (Tyrosinases, Catechol oxidases and Laccases). 100 μL were added to each well right before reading the absorbance every 60 seconds, for 45 minutes at 495 nm. 50 μL of Tris buffer were added instead of samples, as a negative control. The specific PO activity (U/mg protein) was calculated according to the following formula:

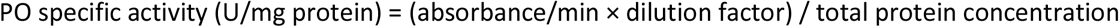

#### Glutathione S-transferase (GST) activity

A volume of 20 μL of visceral mass supernatant at 300 μg/mL was added to 150 μL of phosphate buffer (100 mM PB, pH 6.5) in a 96-well microplate. A mixture (17:1) of reduced *L*-glutathione (GSH, 2.225 mM in PB) and 1-Chloro 2,4 Dinitrobenzene (CDNB, 38 mM in ethanol) was prepared.

180 μL of the GSH/CDNB mixture were added to each well right before reading the absorbance every 60 seconds, for 10 minutes at 340 nm. 20 μL of Tris buffer were added instead of samples, as a negative control.

The specific GST activity (nmol/min/mg of protein) was calculated according to the following formula:

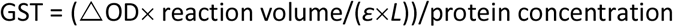

Where ΔOD: time slope of optical density (absorbance/min)

*ε*: molar absorption coefficient of conjugate CDNB = 9.6 mM^−1^cm^−1^

*L*: optical path length through the reaction volume inside the well (cm)

#### Acetylcholine esterase (AchE) activity

A volume of 10 μL of visceral mass supernatant at 300 μg/mL was added to a 96-well microplate with 165 μL of PB (0.1 M, pH 7.8). A solution of dithiobisnitrobenzoate (DTNB, 0.0076 M in phosphate buffer) was prepared. A volume of 10 μL of DNTB was added to each well. A solution of Acetylthiocholine (ATCI, 0.076 M, in water) was prepared. A volume of 5 μL of ATCI was added to each well right before reading the absorbance every 60 seconds, for 10 minutes at 405 nm. 10 μL of Tris buffer were added instead of samples, as a negative control.

The specific AchE activity (nmol/min/mg of protein) was calculated according to the following formula:

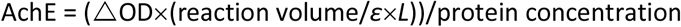

Where ΔOD: time slope of optical density (absorbance/min)

*ε*: molar absorption coefficient of conjugate CDNB = 0.0136 mM^−1^cm^−1^

*L*: optical path length through the reaction volume inside the well (cm)

### Transcriptomic analyses

Real-time PCR was performed to assess differential gene expression in gills and visceral mass of clams. A SV Total RNA Isolation System kit (Promega) was used to extract total mRNA, according to the manufacturer’s instructions. The mRNA concentration was measured in a Take3 plate with a microplate spectrometer (BioTek EPOCH). The RNA purity was checked by comparing the ratio between values measured at 260 and 280 nm and ensured when the ratio was over 2. The RNA samples were diluted to a concentration of 100 ng/μL. They were used for the reverse transcription (RT), using the GoScript Revers Transcription System kit (Promega). RT was performed in an Eppendorf Mastercycler to synthetize cDNAs. Samples were kept at −20°C until their use for qPCR amplifications.

Quantitative polymerase chain reaction (qPCR) were performed in a LightCycler 480 (Roche) using the GoTaq® qPCR Master Mix kit (Promega) in 384-well plates. A mix of primer pair was prepared for each gene (5 μM, Table 1). 1 μL of the pair mix, 12.5 μL of the qPCR Master Mix mixed with 6.5 μL of RNA free water, and 5 μL of cDNA were added to each well. The qPCR program started by heating samples at 95°C for 10 min and then ran 40 cycles as follows: 95°C for 30 s, 60°C for 30 s, and 72°C for 30 s. Samples were gradually heated from 60 to 95°C to get a dissociation curve and check for amplification specificity. Two reference genes were used to normalize gene expression: *β-actin* and *rpl7* (according to the GeNorm method). The differential gene expression was calculated using the 2^−ΔΔCt^ method described by Livak and Schmittgen ^25^. The gene expression was expressed as induction factors compared to controls, according to the following formula:

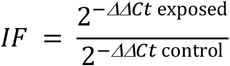

Nineteen genes involved in endocytosis, detoxication, respiratory chain, DNA repair, oxidative stress, apoptosis, neurotransmission and reproduction were assessed (Table 2).

**Table 2:**
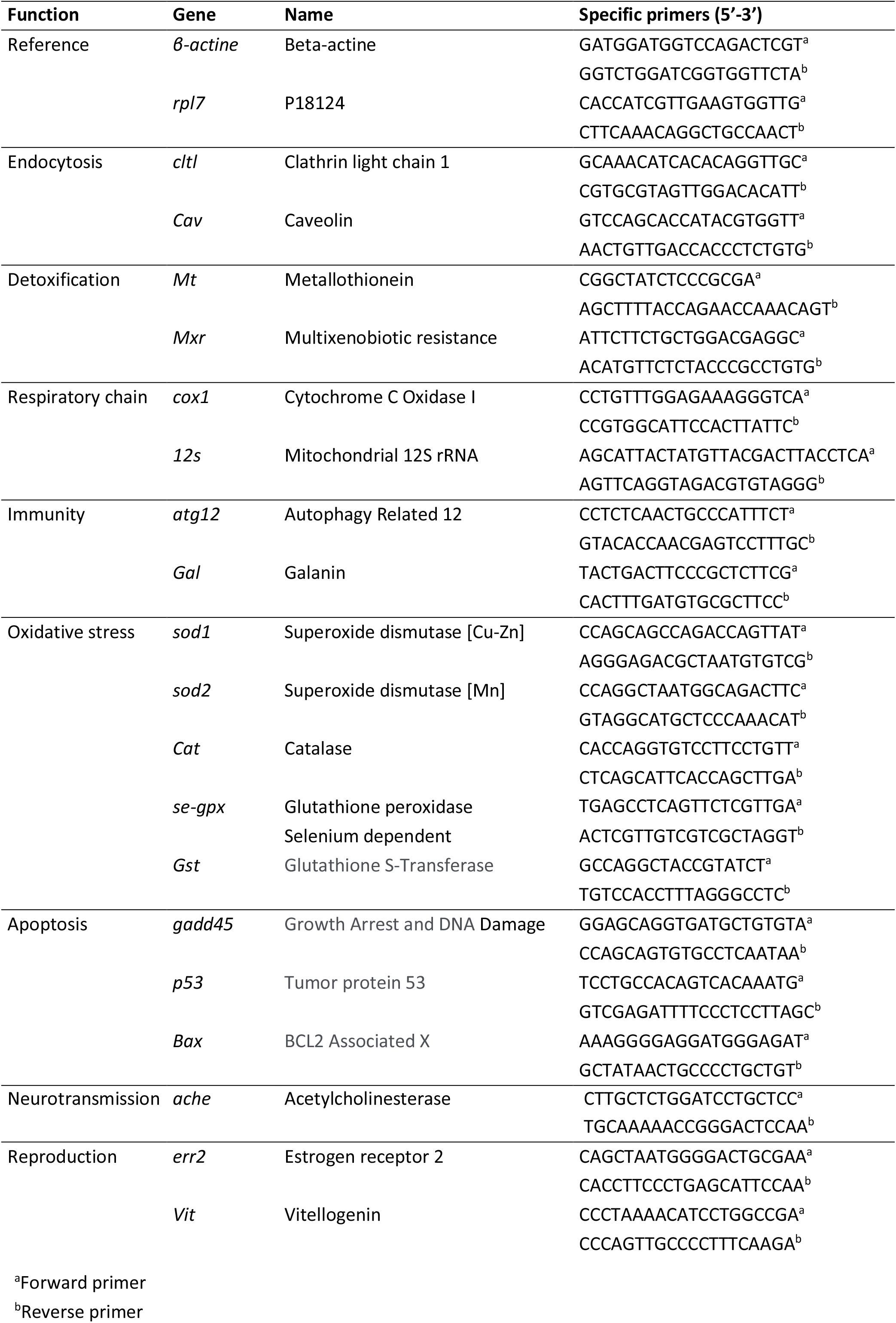
Gene functions studied and primer sequences used

### Data treatment

Significant differences between treatments were tested using non-parametric Kruskal-Wallis tests, with post hoc exact permutation tests for inter-groups comparison after sequential Bonferroni correction (XL-Stat software version 2013.5.09, 1995e2013 Addinsoft). A statistical parameter p < 0.05 was considered significant for all statistical tests.

## RESULTS

### Nanoplastic suspension characterizations

As seen on Figure 1, the crushed environmental nanoplastic sample NP-L contains mostly the classical PE and PP polyolefins. The band near 1700 cm^−1^ indicates the presence also of ester bonds, which might arise either from other common polymers (*eg* PMMA or PET) or be ascribed to polyolefin oxidation into carboxylic functions generated by the outdoor weathering ^26^, in accordance with the negative zeta potential at −32 mV (Table 1). The zeta potential measured for NP-PS that is even more negative than the environmental NP-L might be explained by the adsorption of bicarbonate anions onto hydrophobic surfaces, as shown by Yan et al^27^. In practice, thanks to their negative surface charges, both suspensions of NP-L and NP-PS remained in a dispersed state for several months, as evidenced by the kinetic study by DLS (Figure S2). Right after their preparation, NP-L suspension exhibits a single peak at 334 nm hydrodynamic diameter using the Pade-Laplace multimodal relaxation fitting of the DLS auto-correlogram acquired in backscattering (165° scattering angle), whereas NP-PS show two peaks at 155 nm and 406 nm, with respective weighting factors to the scattered intensity of 57% and 39%. After 3 months, both suspensions still have mean hydrodynamic sizes around 300 nm with polydispersity indexes PDI=0.27-0.28. The slight aggregation at longer time (8 months) could simply be suppressed by sonicating the stock suspensions for 10 min before their usage to contaminate the algae and the clams.

### Al bioaccumulation

Al concentrations were measured in dry weights (DW) of gills and visceral mass of bivalves exposed to both NP-L and Al, or Al alone (Figure 2). Surprisingly, no significant bioaccumulation of Al could be observed in gills, compared to controls, except for the 1 μg/L condition at T7 (203.4 ± 14.8 μg/g DW). Overall, Al concentrations were around 10 times lower in the visceral mass than in gills. The Al bioaccumulation significantly increased in the visceral mass of bivalves exposed to Al alone, at each sampling time, as well as in the Al+ NP-L condition at 10 μg/L and T21 (10,6 ± 1.6 μg/g DW). After 7 days of depuration, there are still significant accumulations in visceral mass of oysters exposed to Al + NP-L 1 and Al.

**Figure 2:**
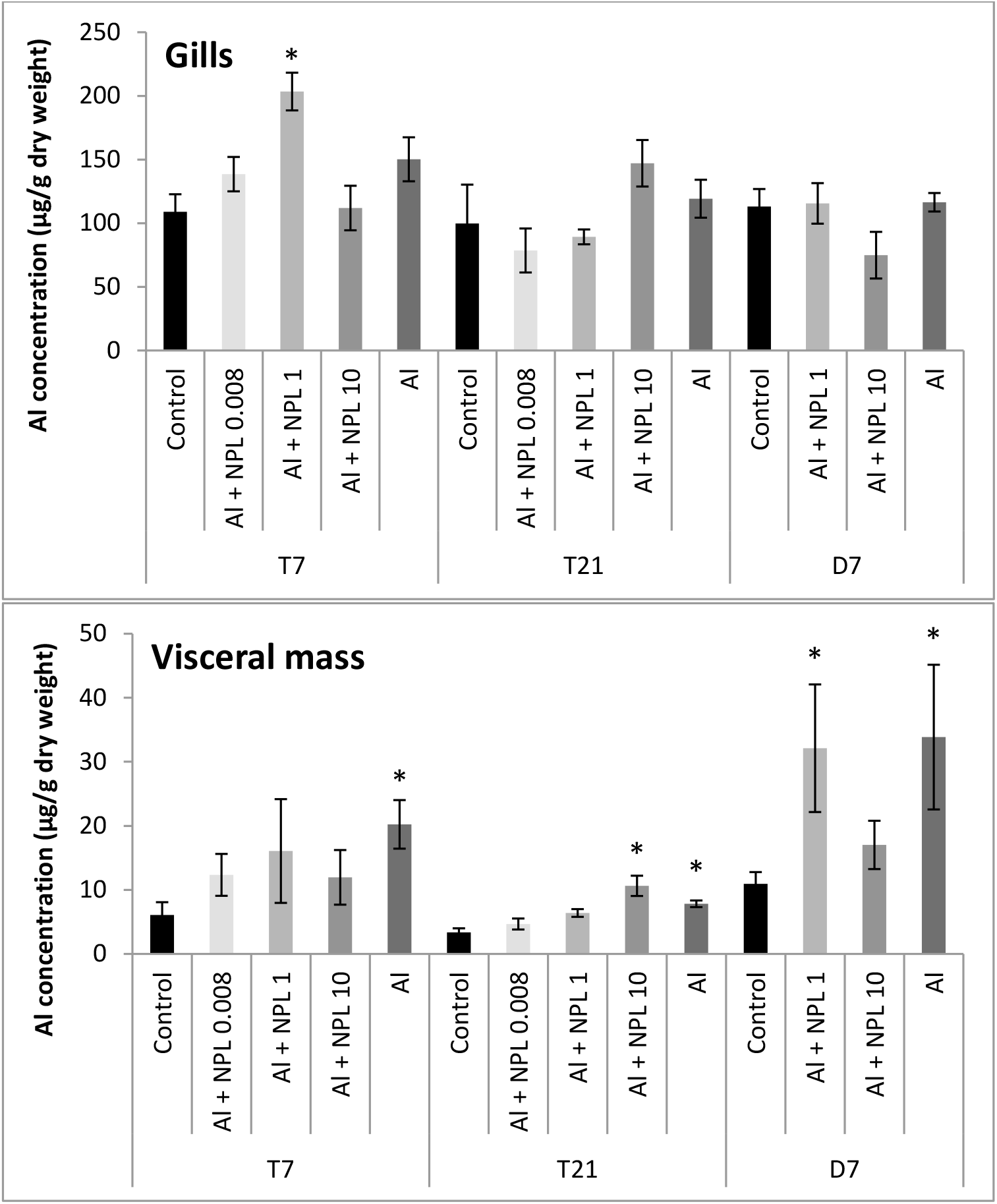
Al concentrations in gills and visceral mass of clams exposed to Al + NP-L or Al alone (mean ± SE, n=4). *: significantly different from controls (p<0.005).

### Transcriptomic analyses

Globally, the gene expressions were more modulated in the visceral mass than in gills (38.8 and 23.7% of genes modulated in the 10 μg/L conditions, respectively), even for the direct exposure to NP-L (Tables 3 and 4). They followed a dose-response trend towards the three concentrations tested. Gene modulations were observed at concentrations as low as 0.008 μg/L in both gills and visceral mass, after direct or dietary exposures. Gene modulations followed the same pattern between the water- and diet borne exposures. However, genes were more up-regulated in gills of clams exposed via the trophic route than the direct route (21 and 16%, respectively for the 10 μg/L conditions after 21 days), whereas the direct exposure triggered more gene modulation in the visceral mass than the trophic route (42 and 26% of genes, respectively for the 10 μg/L conditions after 21 days). Several genes were still up-regulated after 7 days of depuration in the visceral mass and gills of clams exposed through the direct and trophic routes. At 1 μg/L, the waterborne exposure to NP-L modulated more genes than NP-PS in gills (10.5 and 5.3 % of genes up-regulated at T7, respectively), whereas NP-PS triggered more gene up-regulations at 10 μg/L in gills than the waterborne exposure to NP-L (10.5 and 5.3 % of genes up-regulated at T7, respectively). The opposite trend was observed in the visceral mass. Effects of Al were stronger than Al+NP-L in all conditions and times tested, except in gills at 10 μg/L for T7 and T21. Similarly, effects of the waterborne exposure were stronger than the synergic exposure with Al (Al+NP-L) in all conditions tested, except in gills at 1 μg/L for T7 and T21. Effects of Al+NP-L lasted longer than Al and NP-L during the depuration phase.

**Table 3:**
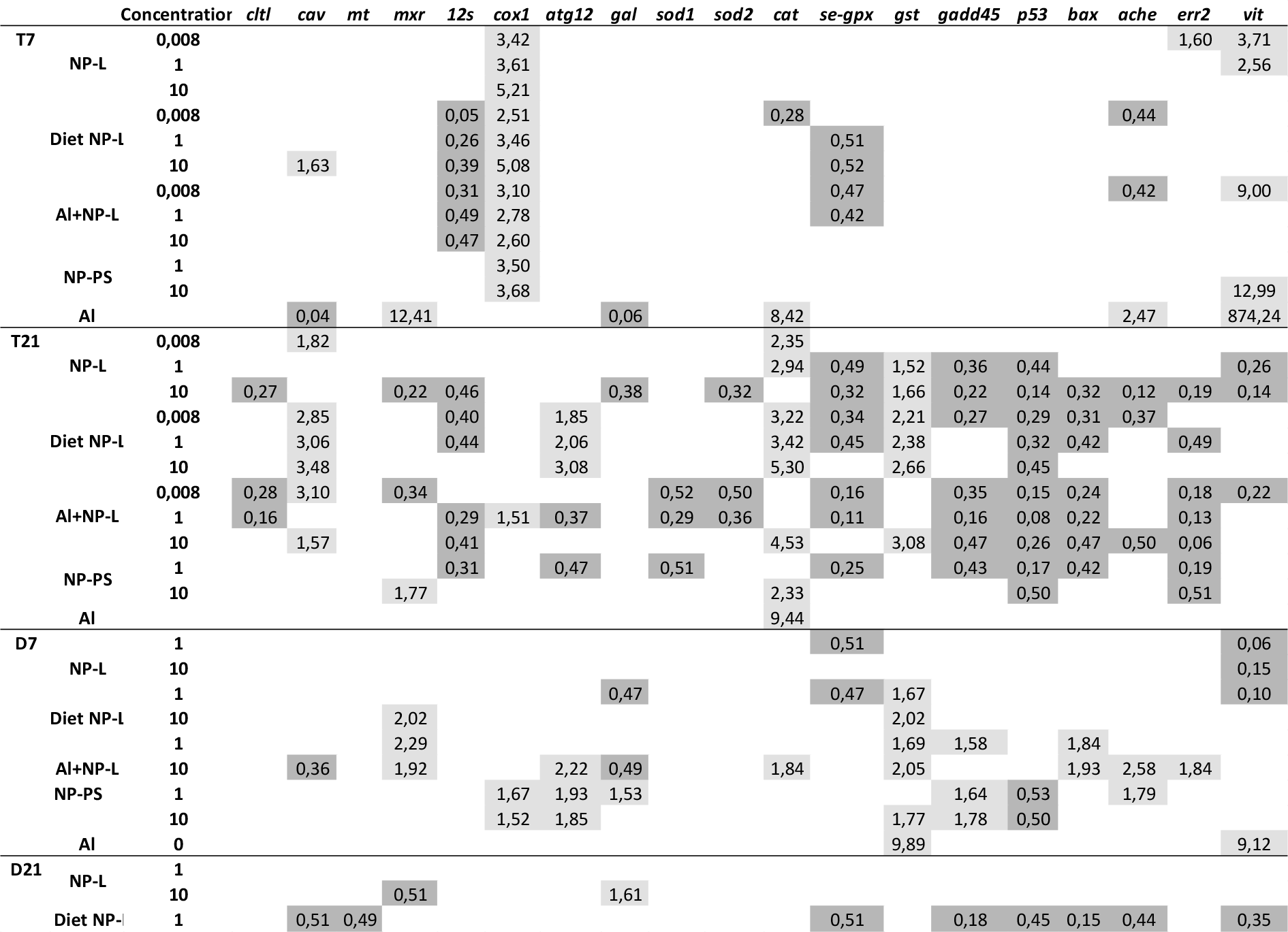
Differential gene expressions observed in gills of clams exposed to the different conditions. Only statistically significant results are reported. Results are given as induction (>1.5) or repression (<0.5) factors as compared to controls.

**Table 4:**
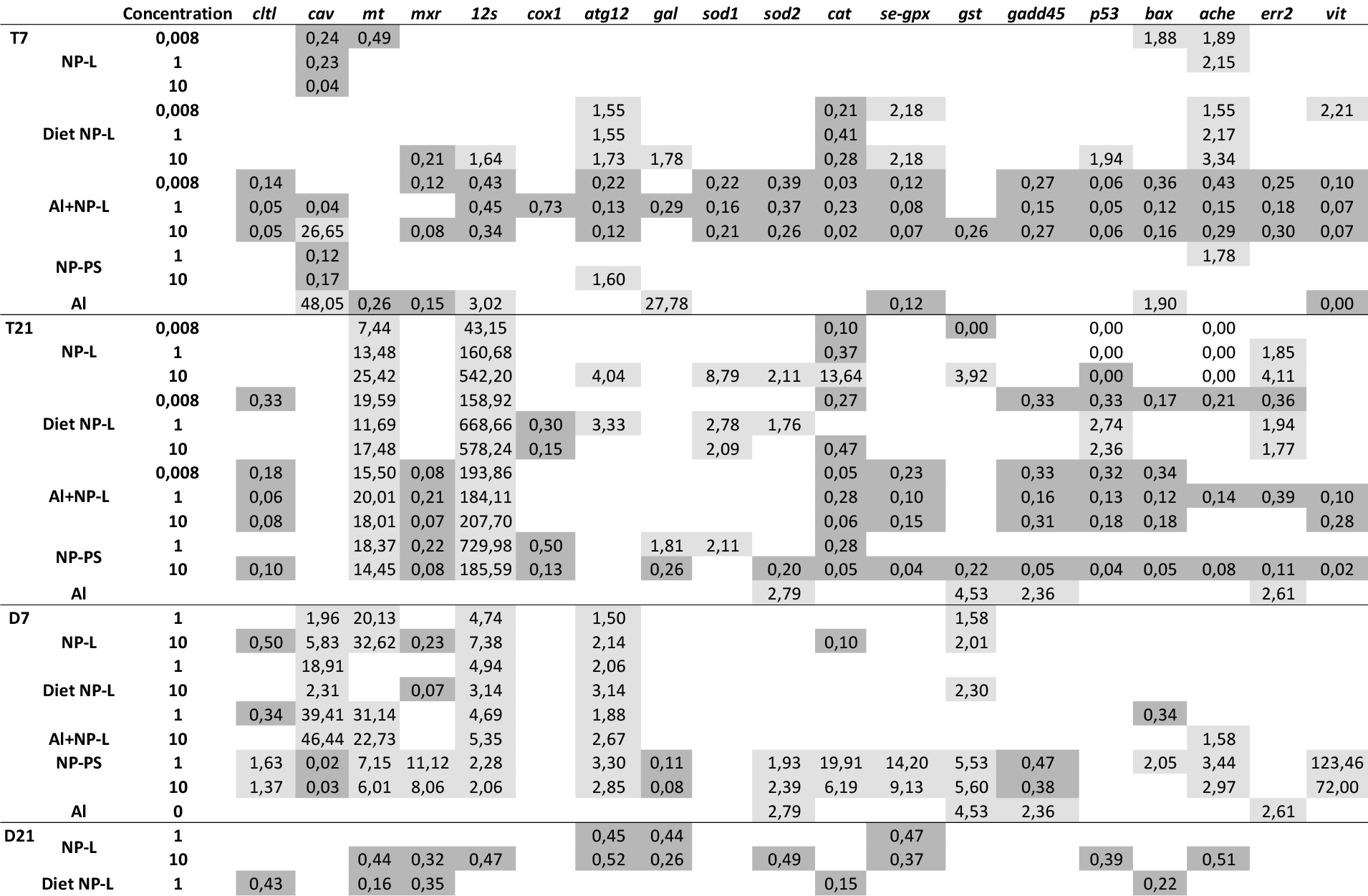
Differential gene expressions observed in visceral mass of clams exposed to the different conditions. Only statistically significant results are reported. Results are given as induction (>1.5) or repression (<0.5) factors as compared to controls.

#### Gills

Genes involved in endocytosis patterns were mainly triggered after 21 days of exposure. No induction of *cltl* was observed, whereas caveolin was up-regulated in the NP-L, Al+NP-L and Diet NP-L conditions (2.85- to 3.48-fold change, for the Diet NP-L condition). Its expression was then repressed during the depuration phase.

No induction of the detoxification gene *mt* was observed in any of the tested conditions, including Al, whereas it led to an increase of the *mxr* expression at T7 (12.41). The *mxr* gene was repressed in NP-L and Al+NP-L conditions at T21 and over-expressed in the NP-PS condition (1.77 for 10 μg/L). It was up-regulated after 7 days of depuration in the Al+NP-L and Diet NP-L conditions and down-regulated after 21 days in the NP-L condition.

The *12s* gene was repressed for the three concentrations of Al+NP-L (0.31-0.47) and Diet NP-L (0.05-0.39) after 7 days. It was also repressed in all conditions except Al after 21 days of contamination. No gene modulation was observed during the depuration phase. On an opposite trend, the *cox1* expression increased in all conditions tested at T7 (2.51 to 5.21), except Al, and in the Al+NP-L condition at T21 (1.51 for 1μg/L). The over-expression lasted until 21 days of depuration in the NP-PS condition.

Low immunity response was triggered during the exposure phase. Only the 3 concentrations tested in the Diet NP-L condition led to an increase of the *atg12* gene (1.85 to 3.08). However, some gene inductions were observed during the depuration phase, namely *atg12* in the Al+NP-L condition (2.22 for 10 μg/L, at D7), *atg12* and *gal* in the NP-PS condition (1.93 and 1.53 for 1 μg/L at D7), and in the NP-L condition (1.61 for 10 μg/L at D21).

Few effects were observed in response against oxidative stress after 7 days of contamination. Only the Al condition led to an up-regulation of the *cat* gene (8.42) whereas *se-gpx* was repressed in the Al+NP-L and Diet NP-L conditions. After 21 days of contamination, *sod1, sod2* and *se-gpx* were repressed in several conditions, namely NP-L, Al+NP-L, NP-PS and Diet NP-L (only for *se-gpx*), whereas *cat* and *gst* were up-regulated in many conditions. During the first 7 days of the depuration phase, *gst* was still up-regulated in all conditions except NP-L.

The genes *gadd45, p53* and *bax* were repressed after 21 days of contamination in all conditions tested except Al. During the depuration phase, concomitant with the induction of *gst*, the exposure to Al+NP-L led to an up-regulation of *gadd45* (1.58 for 1 μg/L) and *bax* (1.84-1.93 for 1 and 10 μg/L, respectively). The NP-PS exposure also led to an increase of *gadd45* expression at D7 (1.64-1.78 for 1 and 10 μg/L, respectively), while *p53* was repressed (0.53-0.50 for 1 and 10 μg/L, respectively).

The neurotransmission gene of AchE was repressed after 7 days in the Al+NP-L and Diet NP-L conditions (0.42 and 0.44 for 0.008 μg/L, respectively), while it was over-expressed in the Al condition (2.47). The gene *ache* was repressed after 21 days of exposure in the conditions Al+NP-L (0.50 for 10μg/L), NP-L (0.12 for 10 μg/L), and Diet NP-L (0.37 for 0.008 μg/L). However, its expression was up-regulated during the 7 first days of depuration in the Al+NP-L, Diet NP-L and NP-PS conditions (2.58, 1.49 and 1.79, respectively).

The reproduction pathways were also affected after 7 days of contamination, through the direct route with the up-regulation of the *err2* (1.39-1.60) and *vit* genes (2.56-3.71). All conditions led to an up-regulation of *vit* after 7 days of exposure, except the Diet NP-L condition. However, *err2* was repressed after 21 days of exposure in all conditions except Al. NP-L and Al+NP-L conditions also led to a down-regulation of the *vit* gene. During the depuration phase, the Al+NP-L and Al conditions triggered an over-expression of *err2* and *vit* (1.84 for 10 μg/L, and 9.12, respectively), whereas the water- and diet borne exposures led to a down-regulation of *vit*, which lasted until D21 for the Diet NP-L condition.

#### Visceral mass

The endocytosis pathways were poorly induced during the exposure phase. Both *cltl* and *cav* genes were repressed after 7 and 21 days of contamination, except in the Al+NP-L and Al conditions which led to an over-expression of *cav* (26.65 for 10 μg/L and 48.05, respectively). However strong over-expressions were observed for *cav* during the 7 first days of depuration in most of the conditions except Al and NP-PS (up to 46.44-fold change for Al+NP-L at 10 μg/L). The *cltl* gene was up-regulated in the NP-PS condition after 7 days of depuration, whereas *cav* was down-regulated. The *mt* gene was strongly up-regulated, except for the Al condition, after 21 days of contamination (from 7.44 to 25.42 fold change for the NP-L condition, for instance). The up-regulation lasted until 7 days of depuration, and then the *mt* expression was repressed. The *mxr* expression was down-regulated throughout the experiment, in most of the conditions tested. Only NP-PS led to an up-regulation of *mxr* after 7 days of depuration.

The mitochondrial metabolism was affected with a strong increase of the *12s* gene expression after 21 days of contamination, in all conditions tested. The induction reached 542.20 and 578.24-fold changes in the NP-L and Diet NP-L conditions at 10 μg/L. The over-expression was observed until 7 days of depuration, except for the Al condition, and was then repressed. The *cox1* gene was down-regulated after 21 days of exposure through the trophic route and to NP-PS.

The immunity was also strongly induced, through the *atg12* gene. It was up-regulated after Diet NP-L exposure from T7 to D7. The NP-L exposure also led to an increase of *atg12* expression (3.33 for 10 μg/L). The gene up-regulation was observed until 7 days of depuration in all conditions tested, except Al, and was then repressed. The *gal* gene was strongly induced after 7 days of exposure to Al (27.78) and 21 days to NP-PS (1.81 for 1 μg/L) but not for other conditions.

The exposure to Al+NP-L led to a strong down-regulation of all genes involved against oxidative stress after 7 days and of *cat* and *se-gpx* after 21 days. The exposure to Diet NP-L led to a down-regulation of *cat* and an increase of *se-gpx* expression (2.18 for both 0.008 and 10 μg/L) after 7 days. No effects were observed for the NP-L and NP-PS conditions after 7 days of exposure. After 21 days of exposure, *cat* and *se-gpx* expression were still repressed in the Al+NP-L condition, whereas the Al condition up-regulated *cat* and *gst* expressions (1.52 and 4.83, respectively). An induction of *sod1, sod2, cat* and *gst* was observed for the highest concentration tested for the NP-L condition (8.79, 2.11, 13.64, 3.92, respectively). The Diet NP-L exposure also led to the up-regulation of *sod1, sod2* and *cat* whereas the NP-PS exposure decreased the expression of *sod2, cat, se-gpx* and *gst*.

During the depuration phase, the *gst* gene was induced after 7 days, in all conditions tested except Al+NP-L. The NP-PS exposure also triggered the up-regulation of *sod2, cat, se-gpx* after 7 days of depuration.

The *gadd45* gene involved in DNA repair was down-regulated after 7 and 21 days of exposure in several conditions, especially the Al+NP-L. A low apoptotic response was observed, with the down-regulation of *p53* and *bax* after 7 days of exposure to Al+NP-L and in most of the conditions tested, after 21 days. Only the Diet NP-L condition led to an over-expression of *p53* (1.94 and 2.36 for 10 μg/L after 7 and 21 days, respectively) while *bax* was up-regulated after 7 days of exposure to Al and NP-L (1.90 and 1.88 for 0.008 μg/L, respectively). No clear trend was observed towards the apoptotic response during the depuration phase.

The neurotransmission was assessed through the *ache* expression, which was up-regulated after 7 days of exposure through the direct and trophic routes, as well as NP-PS, whereas it was down-regulated in the Al+NP-L condition. After 21 days, the *ache* expression was repressed in all conditions, except Al. During the depuration phase, its expression was again over-expressed in the Al+NP-L and NP-PS conditions.

The reproduction pathway was also affected by several conditions, especially after 21 days. The direct and trophic routes triggered an increase in *err2* expression (1.85-4.11 and 1.94-1.77 for the 1 and 10 μg/L, respectively, for direct and trophic conditions, respectively). Its expression was decreased in the Al+NP-L condition after 7 and 21 days, as well as the *vit* expression. The expression of *err2* and *vit* was decreased in the NP-PS condition after 21 days, and increased after 7 days of depuration (123.46 and 72 for *vit* at 1 and 10 μg/L respectively).

### Filtration test

The filtration test was performed after 21 days of experiment (Figure 3). The exposure to NP-L led to a significant decrease of the filtration rate after 60 min in the 1 μg/L condition (129900 ± 27600 cells/ml/individual, compared to control reaching 336800 ± 75200 cells/mL/individual). The bivalves exposed to Diet NP-L showed an opposite behaviour, with a significant increase of their filtration rate after 30 and 45 min, for the 1 and 10 μg/L concentrations (345000 ± 47070 and 328600 ± 24200 cells/mL/individual, respectively after 45 min, compared to control reaching 195450 ± 35200 cells/mL/individual). It was the same observation in the NP-PS condition, with increasing filtration rates in clams exposed to 1 and 10 μg/L, but only in the first times of the test (5 to 20 minutes). The filtration rates averaged 162000 ± 36000 and 248000 ± 88000 cells/ml/individual for the 1 and 10 μg/L after 20 min, while the control did not go higher than 104 000 ± 48500 cells/mL/individual. No significant results were obtained for the Al+NP-L condition.

**Figure 3:**
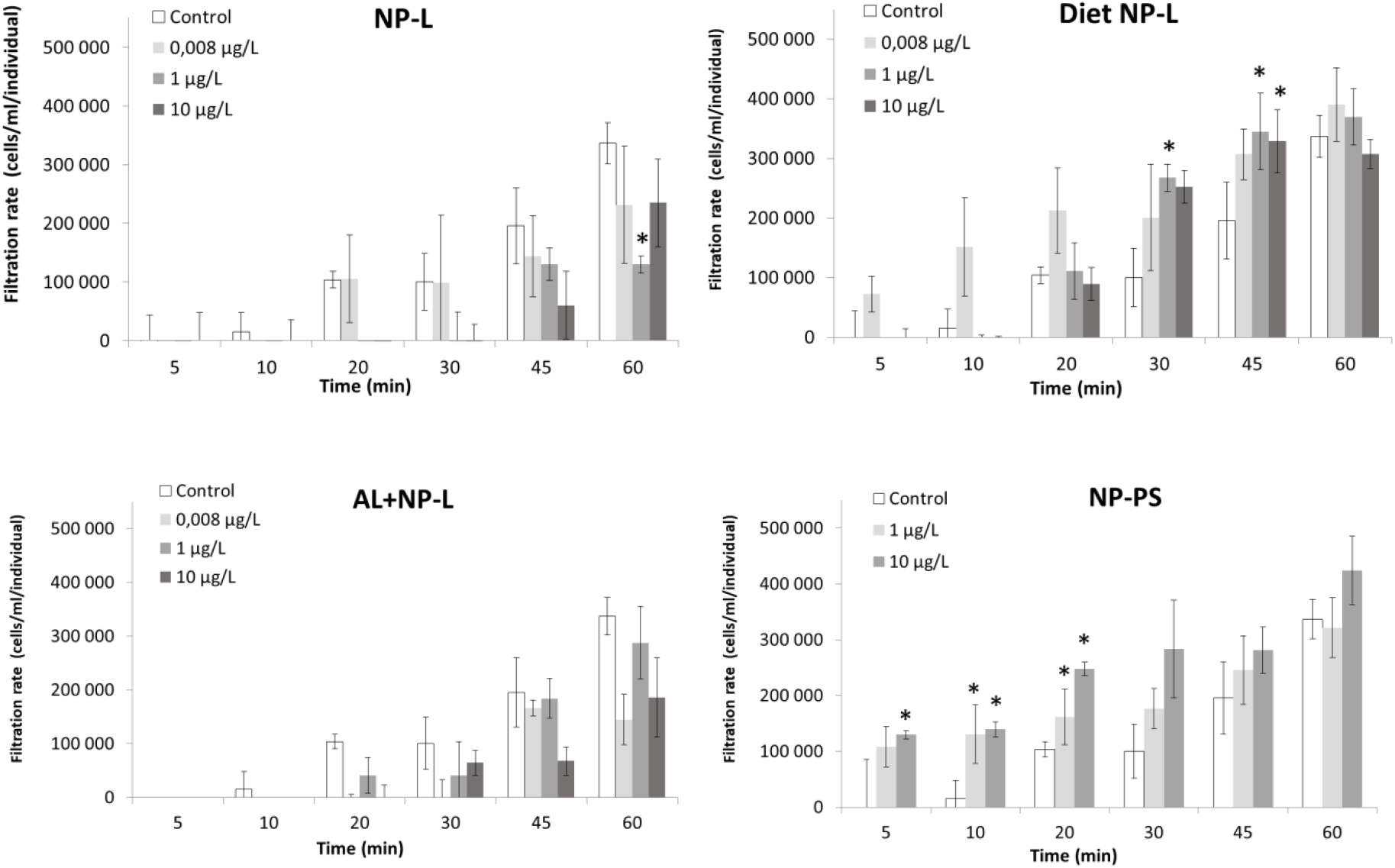
Filtration rates of clams exposed to the different conditions (mean ± SD, *n*=4). *: significantly different from controls (Student test *p*<0.005).

### Phenol oxidase activity

The phenol oxidase (PO) activity was measured after 7 and 21 days of exposure and after 7 days of depuration (Figure 4). Very few results were different from controls. After 7 days of exposure to NP-PS, the activity was significantly higher than in controls, reaching 0.68 ± 0.04 and 0.32 ± 0.08 U/mg of proteins, respectively. No other significant result was observed after 7 and 21 days of exposure. After 7 days of depuration, the clams from the Diet NP-L condition at 10 μg/L showed an activity significantly higher than controls (0.83 ± 0.08 and 0.36 ± 0.01 U/mg of proteins, respectively), but not the other conditions.

**Figure 4:**
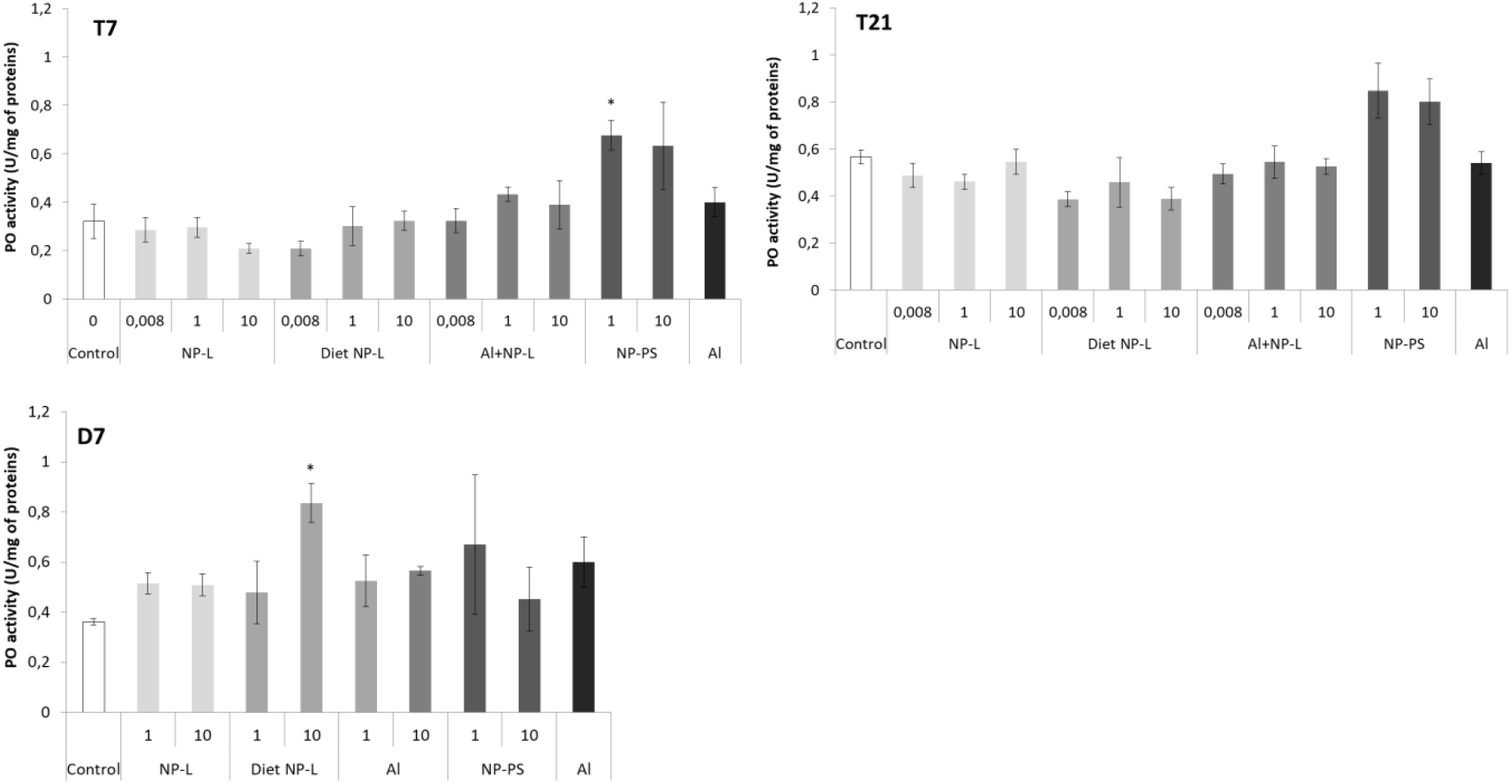
Phenol oxidase activity in gills of clams exposed to the different conditions (mean ± SD, *n*=4). *: significantly different from controls (Student test coefficient *p*<0.005).

### GST activity

No difference in the GST activity was observed after 7 days of contamination (Figure 5). After 21 days, the GST activity was significantly lower in the Al+NP-L condition at 1 μg/L compared to controls (1.60 ± 0.26 and 2.97 ± 0.26 nmol/min/mg of proteins, respectively). After 7 days of depuration, the activity was also significantly decreased in the NP-L and Diet NP-L conditions, at 1 μg/L, compared to controls (2.07 ± 0.25, 1.78 ± 0.26 and 2.80 ± 0.14 nmol/min/mg of proteins, respectively).

**Figure 5:**
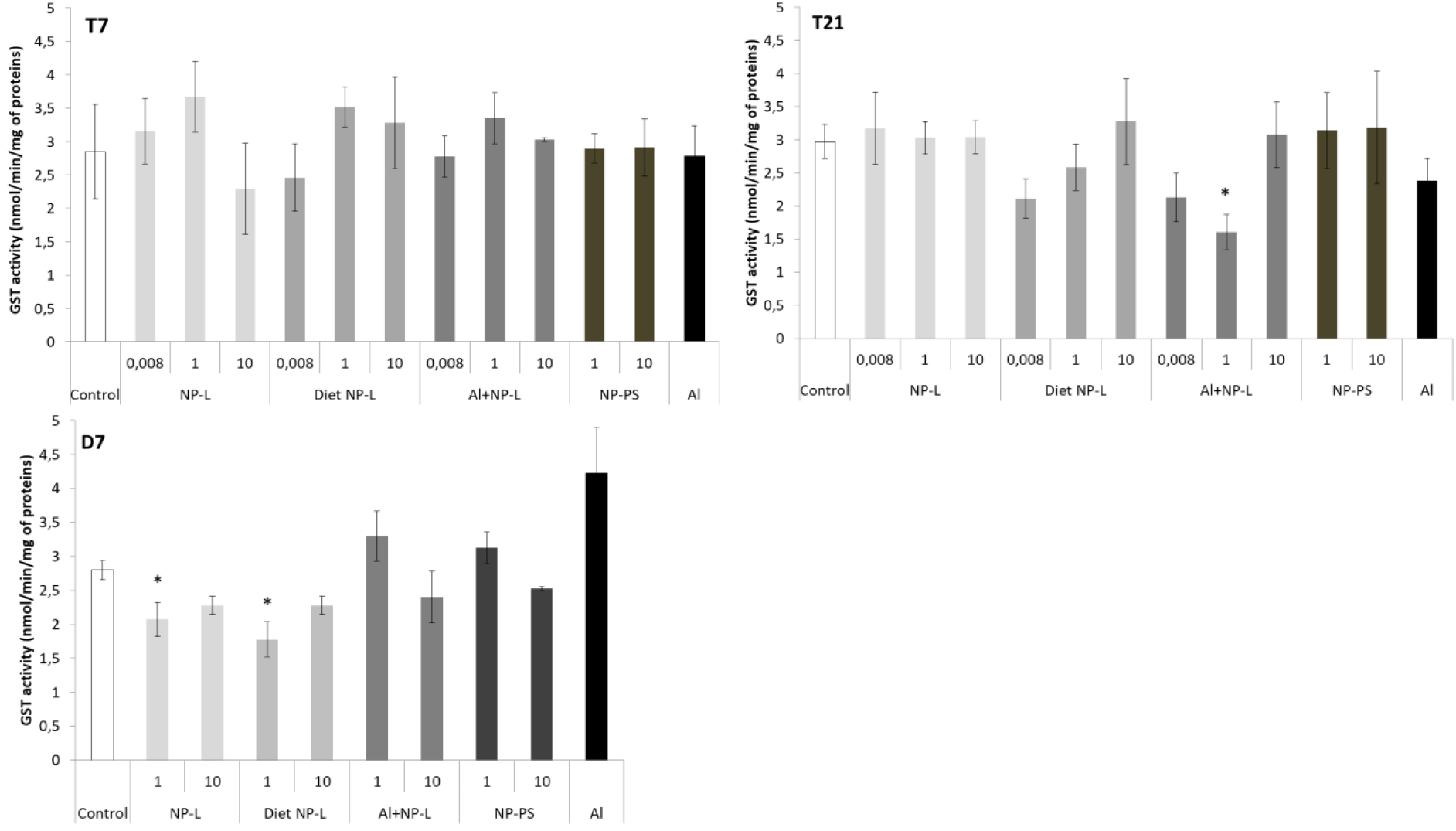
GST activity in the visceral mass of clams exposed to the different conditions (mean ± SD, *n*=4). *: significantly different from controls (*p*<0.005).

**Figure 6:**
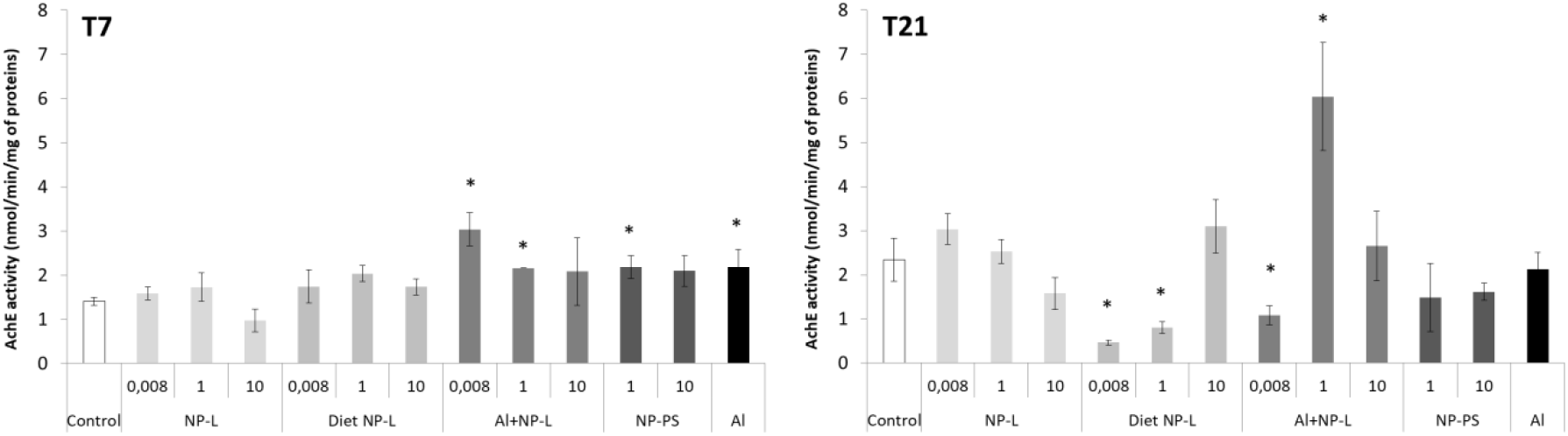
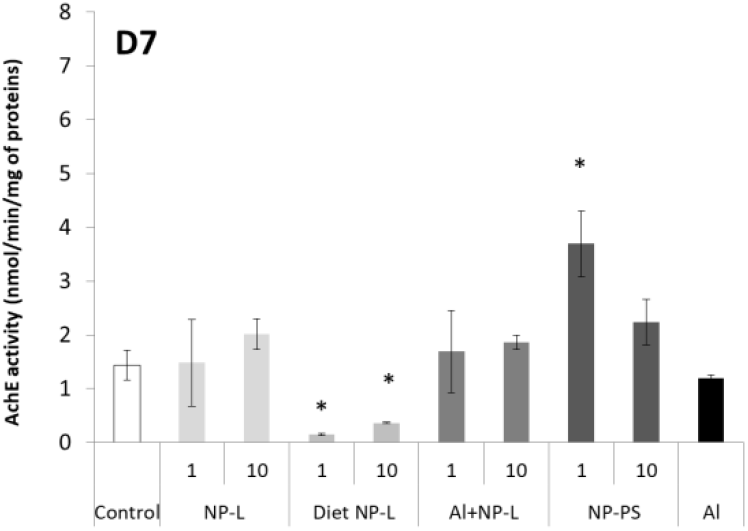
AchE activity in visceral mass of clams exposed to the different conditions (mean ± SD, *n*=4). *: significantly different from controls (*p*<0.005).

### AchE activity

After 7 days, the activity of AchE was significantly higher in bivalves exposed to Al+NP-L at 0.008 and 1 μg/L (3.04 ± 0.37, 2.16 ± 0.02 nmol/min/mg of proteins, respectively) and to NP-PS at 1 μg/L and Al (2.18 ± 0.25 and 2.18 ± 0.38 nmol/min/mg of proteins, respectively) than controls (1.40 ± 0.09 nmol/min/mg of proteins). After 21 days, the activity of AchE significantly decreased in the Diet NP-L condition (0.46 ± 0.06 and 0.80 ± 0.13 nmol/min/mg of proteins for 0.008 and 1 μg/L of Diet NP-L, respectively) compared to controls (2.34 ± 0.49 nmol/min/mg of proteins), as well as in the Al+NP-L condition at 0.008 μg/L, on the contrary to an increase at 1 μg/L. Similarly, the activity decreased after 7 days of depuration in the Diet NP-L condition for the 1 and 10 μg/L concentrations, and increased in the NP-PS condition at 1 μg/L.

## DISCUSSION

### Aluminium bioaccumulation

No significant bioaccumulation of Al could be observed in gills of bivalves exposed to either Al or Al + NP-L, except for one condition: Al+NP-L at 1 μg/L and 7 days. Rosseland et al, ^28^ demonstrated that Al could provoke an osmoregulatory failure in gills and led to the precipitation of Al. This was possibly due to an acidification of mucus secretion that prevented from Al binding to gill cells.^29^.

Surprisingly, Al concentrations were lower in the visceral mass compared to gills in controls. However significant increases were observed in the bivalves exposed to Al alone or punctually in conditions Al+NP-L. It suggests that NPs may affect the metal accumulation. It could result from a lower bioavailability of Al in presence of NPs, potentially due to Al adsorption onto NPs’ surface. It has been suggested that NPs can be directly rejected through pseudo-faeces or mucus secretion ^30, 31^ and that the bioavailability of a metalloid such as arsenic was lowered in presence of NPs in mangrove oysters, due to its adsorption to the surface of NPs ^15^. On the assumption that a part of Al is bound to NPs, it could explain, at least partly, the low accumulation of Al in gills and visceral mass in conditions with NPs.

### Transcriptomic analyses

Our study aimed to compare gene expression after water- and diet borne exposures to NPs, with NP-L or NP-PS commonly used in the literature, as well as synergic effects with a metal: aluminium. Two organs were investigated to see differences between exposure routes: gills and visceral mass.

#### Differences between gills and visceral mass

Gene modulations were observed at concentrations as low as 0.008 μg/L in both gills and visceral mass, after direct or dietary exposures. Globally the gene expressions were more modulated in the visceral mass than in gills (38.8 and 23.7% of genes modulated in the 10 μg/L conditions, respectively), even for the direct exposure. Similarly, a study on *Corbicula fluminea* exposed to NP-PS (80 nm, 0.1-5 mg/L for 96h) showed that the visceral mass, as the main organ involved into metabolism and detoxification, presented a stronger response against oxidative stress than gills ^17^.

Moreover, the cell permeability of gills, in direct contact with the surrounding medium, can act as a protective barrier against xenobiotics. This could reduce pollutant accumulation through the direct route ^32^. We observed through the waterborne exposure a strong production of mucus coming out from the clams’ valves which could also act as a protective mechanism of gills towards contamination by trapping NPs and rejecting them into the surrounding medium. This phenomenon has also been reported on oysters exposed to NPs ^31, 33^.

After 21 days of exposure, gills were mainly responding to the surrounding contamination through the endocytosis pathway, with the up-regulation of the caveolin gene in all conditions tested, except NP-PS. The endocytosis was concomitant with the decrease in the *12s* expression in the NP-L condition and with the up-regulation of *gst* and *cat*. This could underline a disturbance of the energetic balance leading to an oxidative stress in clams after NP-L entry. Similar trends were observed in previous studies on the pacific oyster *I. alatus* exposed through the direct route to NPs, with the up-regulation of *cav, 12S* and *cat* genes ^33^, whereas it was not observed after dietary exposure ^15^. Surprisingly, no response was observed towards apoptosis in gills, with a global inhibition of *gadd45, p53* and *bax* in all conditions tested (except Al). It suggests that responses against oxidative stress in gills could have efficiently counteracted the effects of NPs and did not end up in apoptosis.

Results in visceral mass were more pronounced than in gills after 21 days of exposure. We observed a very strong induction of the *mt* (except in the Al condition) and *12s* genes in all conditions tested, while *cox1* was repressed in the Diet NP-L and NP-PS conditions. The results suggest that metallothioneins, well known for their key role in metal detoxification ^34^, could also be involved in the case of NP exposure, probably through the oxidative stress generated by their high surface reactivity. They could have also react to metals or additives released in the tissues after absorption of NPs in the cells to sequester or excrete them. The fact that Al did not trigger an up-regulation of *mt* was already described in the literature in freshwater molluscs such as *Dreissena polymorpha* ^35^. But considering the very low bioaccumulation observed in tissues, the weak concentrations of Al may also have not been sufficient to implement a response towards the *mt* gene, even if this metal generated an oxidative stress (*cat* induction). This comforts our hypothesis that the *mt* gene can actually be modulated by NPs themselves, and not only in case of combined exposure with metals. To our knowledge, this is the first evidence of *mt* involved in the response against NPs in bivalves. Unlike in gills, we observed in the visceral mass a very strong increase of *12s* expression in all conditions tested. Such up-regulations have been reported in other studies on the oyster *I. alatus* after a direct exposure to environmental NPs for one week ^31, 33^. This may show a trial to increase mitochondria number that could be at least partly related to the increase in energy demand to excrete NPs. Again, Al alone led to much lower inductions than Al+NP-L, showing that gene responses are mainly driven by NPs themselves.

We also observed an increase of the expressions of genes involved in immunity, oxidative stress and apoptosis in visceral mass (*atg12, sod1, sod2* and *cat*). This led to an apoptotic response in the Diet NP-L condition only, with the up-regulation of *p53*, as shown on the pacific oyster *I. alatus* ^33^.

Genes involved in the reproduction were also tested and results showed an up-regulation of *err2* in the visceral mass of clams exposed through the direct and trophic routes (1 and 10 μg/L), but no effects were observed on the vitellogenin gene. This underlines the first evidence of reproductive gene impairments after an exposure to NPs at such low concentrations. Reproductive adverse effects have already been described in pacific oysters exposed to 50 nm NP-PS but at higher concentrations (0.1, 1, 10 and 25 mg/L). The authors showed that NP could alter fertilization success, lead to larval teratologies, and arrest of growth development ^36^.

#### Differences between water- and diet borne exposures to NPs

Overall, genes were more up-regulated in gills of clams exposed via the diet borne exposure than via the waterborne (21 and 16%, respectively for the 10 μg/L conditions after 21 days), whereas the waterborne exposure triggered more gene modulation in the visceral mass than the diet borne route (42 and 26% of genes, respectively for the 10 μg/L conditions after 21 days).

After 7 days of exposure, very few effects were observed in the waterborne condition, both in gills and visceral mass. The strongest gene modulations were shown after the diet borne exposure in the visceral mass, with the up-regulation of gene expressions among all functions tested (namely *12s, atg12, gal, se-gpx, p53* and *ache*) and in a dose-response manner. This suggests that NPs may trigger more effects on bivalves *via* the diet borne route than the waterborne one. Another study comparing water- and diet borne exposures to nanoparticles of gold in *Corbicula fluminea* showed that the effects on gene expressions were much lower after the diet borne exposure ^37^. The authors suggested that it was probably due to valve closures occurring in response to water contamination. This phenomenon has been documented for metals in *Corbicula fluminea* ^38^ but owing to the lack of analytical methods available to quantify NP in tissues, it is difficult to actually rely toxicity to NP bioaccumulation. However, our results from the filtration test showed that the ventilatory activity of *Corbicula fluminea* decreased after the waterborne exposure, while it increased after the diet borne exposure. This suggests a stronger NP accumulation in the Diet NP-L condition, coinciding with the up-regulation of most genes tested in this condition.

Gene expressions in gills followed the same trends after 21 days of exposure to either the water or the trophic route. The main differences were observed in the visceral mass. Both conditions led to an increase of gene expressions of *mt, 12s* and *atg12*. However, responses differed for genes involved in oxidative stress response and apoptosis. The direct exposure triggered a response against oxidative stress but not towards apoptosis, whereas we observed the opposite pattern for the trophic route in visceral mass.

#### Differences between NP-L and NP-PS

There were few differences on gene expressions after 7 days of contamination to either NP-L or NP-PS, in gills or visceral mass. However, after 21 days, endocytosis pathways were induced in gills of clams exposed to all conditions, except NP-PS. In the visceral mass, most of genes of clams exposed to NP-PS were repressed, except *mt* and *12s*, while they were up-regulated in the NP-L condition for the different concentrations tested. It seems that the cell response towards NP-L exposure occurs faster than with NP-PS. Indeed, NP-PS did not trigger an up-regulation of genes involved against oxidative stress and towards apoptosis until bivalves were maintained under depuration conditions (D7). No such discrepancy between NP types has been shown in other studies so far under saline conditions ^31, 33^. The lack of gene up-regulation during NP-PS exposure underlines how the origin, morphology, oxidation state, and additional xenobiotics can interact with internalization and adverse pathways in clams. It underlines the importance to consider naturally aged environmental NP in ecotoxicological studies instead of reference nanospheres of plastic such as NP-PS. This is in agreement with Arini et al ^31^ who concluded that environmental NPs had more gene effects than NP-PS after 7 days of contamination in the marine oyster *I. alatus*. Being more oxidized than NP-PS, environmental NPs could present more negatively charged groups onto their surface ^39^, which could end up in earlier cell up-take and effects than with NP-PS. It has also been suggested that NPs’ surface charges play a role in damaging mitochondrial function by inducing a direct accumulation inside mitochondria ^40^. Instability in mitochondrial membrane potential can produce ROS and induce an oxidative stress ^41^. This corroborates our results showing earlier response in bivalves exposed to NP-L, compared to NP-PS, namely towards oxidative stress and apoptosis.

#### Differences between NP-L and Al+NP-L

Few differences were observed between NP-L and Al+NP-L responses in gills. However, in the visceral mass, the Al+NP-L exposure led to synergic effects with a decrease of the expression of most of genes tested. It shows how deep the general state of clams was affected by the combined contamination of a metal and NPs. When genes are repressed as described here, it usually demonstrates a loss of fundamental functions leading to death. For instance, some authors demonstrated that cellular transcription can be altered at the genome-wide level after DNA damage ^42^. Similar synergic effects were shown after waterborne exposure to environmental NPs added to arsenic on pacific oysters ^31^.

After 21 days of contamination in the visceral mass, the effects of Al alone led to much lower inductions of *mt* and *12s*, than after the Al+NP-L exposure, showing that responses are mainly driven by NP exposures. Moreover, effects of Al+NP-L lasted longer in the depuration phase than the single Al or NP-L exposures.

#### Depuration

Gene responses were still implemented after 7 days of depuration in gills after the diet borne exposure (*gst, ache*) and in the visceral mass after both water- and diet borne exposures (*cav, mt, 12s, atg12, gst*).

We also observed delayed effects of NP-PS and Al+NP-L with the up-regulation of genes involved in the detoxication, DNA repair, oxidative stress and apoptosis occurring in gills and visceral mass during the depuration phase. This suggests that clams are recovering from the exposure and able to implement gene responses against the remaining stress. After 21 days of depuration, most of the tested genes were repressed in gills and visceral mass. A study on *Pecten maximus* showed that 68% of the initial burden of 250 nm ^14^C-radiolabeled NP-PS had been purged after 3 days of depuration ^43^. This suggests fast depuration capacities in bivalves, which is in agreement with our results on gene up-regulations, lasting no longer than 7 days after the end of exposure.

#### Differences between effects on freshwater and marine species

After 7 days of exposure, there was a significant increase of the *cox1* expression in gills for all conditions, except Al, and a down-regulation of *12s* and *se-gpx* in the Diet NP-L condition. These results are quite different from other studies made under marine conditions on the pacific oysters. The expression of *cox1* in gills of *Isognomon alatus* was not modulated after neither water- nor diet borne exposures to environmental NPs for 7 days ^*15*, 31^. However, the expression of *12s* was up-regulated in gills of *I. alatus* after direct exposure to environmental NPs ^31, 33^ while it was down-regulated in the present study. Overall, similar studies conducted so far under saline conditions showed effects following a reverse dose-response relationship ^33^ whereas in the present study the gene modulation was dose-dependent. Those results point out the importance of considering environmental conditions (namely salinity) and NPs’ composition, degradation status and adsorbed contaminants to investigate NPs’ toxicity on bivalves, as they can act on their aggregation and then bioavailability ^44^.

### Immune and anti-oxidative responses through PO and GST activities

Transcriptomic results showed that NPs triggered an oxidative stress and immune responses. However, very few immune and anti-oxidative responses were observed at the enzymatic level, regarding to PO and GST activities. Some studies have shown an inhibition of GST and PO activities in *Corbicula fluminea* in response to NP-PS exposure for 96h ^17^, but the authors used higher concentrations than in the present study, ranging from 0.1 to 5 mg/L. It is possible that concentrations used in this study were not sufficient to induce an enzymatic response.

### Neurotoxicity through AchE activity and filtration behaviour

#### AchE and gene expression

AchE results showed good relationships between gene expression and enzyme activity. After 21 days of trophic exposure we showed a significant decrease in gene expression and enzyme activity of AchE in the visceral mass. The opposite pattern was shown after 7 days of contamination with NP-PS, with an up-regulation of *ache* leading to an increase of the AchE activity in the visceral mass. However no effects were shown after 21 days of experiment. The relationship between gene expression and enzyme activity was less clear for the NP-L and Al+NP-L exposures.

Inhibitions of AchE following an NP exposure have been reported in several species. For instance, a study on zebrafish reported a decrease in AchE activity after a 72h exposure to 1 mg/L of 50 nm NP-PS ^14^. Another study on *M. galloprovincialis* also resulted in a significant decrease in the cholinesterase activity after a 96h exposure to NP-PS (0.05 up to 50 mg/L) ^45^. It has been suggested that a modification of the AchE activity could result from the ROS production, namely H_2_O_2_, as it could promote disturbances in the cholinergic system by altering AchE structure ^46^. In our study, the AchE activity was significantly decreased after 21 days of diet borne exposure, coinciding with the strongest induction of genes involved against oxidative stress. However no such relation could be observed for other conditions.

#### AchE and filtration behavior

The activity of AchE measured in the visceral mass was significantly inhibited after 21 days of contamination to Diet NP-L and NP-PS. It lasted until 7 days of depuration, suggesting a neurotoxicity resulting from NP exposure. The nerves present into the visceral mass can control the valves opening and closure ^17^. Li et al ^17^ also observed an inhibition of AchE in *Corbicula fluminea* exposed to NP-PS for 96h (0.1 to 5mg/L). It resulted in an increased retention time of fluorescent NPs in organs, leading to a stronger NP accumulation in *Corbicula fluminea*. Valve opening is the main process regulating the filtration rate and thus the contaminant accumulation and excretion. The AchE inhibition in Diet NP-L and NP-PS conditions are concomitant with the results obtained for the filtration test. Indeed, we observed an increase of the ventilatory activity in clams exposed to these conditions, which could clearly be related to the inhibition of AchE. Authors suggested that its inhibition can directly lead to an excess of acetylcholine in the nervous system and could affect several functions such as valve opening and closing ^47^. By disturbing the filtration behaviour of bivalves, NPs could have direct effects on xenobiotic accumulation and excretion capacities. This corroborates our transcriptomic results, showing more effects triggered after the Diet NP-L and NP-PS exposures compared to the NP-L exposure. To our knowledge, no such relationship between AchE activity and filtration rate has been shown so far in freshwater bivalves after exposure to NPs.

A study on *Corbicula fluminea* exposed to environmental NPs (1-10 000 μg/L) did not show any effect on the filtration rates of clams after 48h of direct exposure ^30^. However, the authors showed an increase in pseudo-faeces production, suggesting an effect on excretion. Our results report the first evidence linking an AchE altered activity (gene repression and enzyme inhibition) to actual behavioural impairments in bivalves exposed to NPs.

## CONCLUSION

This study aimed to test different exposure routes, types of NPs and biological endpoints to have an integrative view of NPs’ toxicity on freshwater bivalves. Results have shown that NPs may be responsible of behavioural, enzymatic and gene expression disorders, which may result in direct effects on contaminant accumulation/excretion, and individual growth, reproduction or survival. This study clearly highlighted how differently NPs can affect biological mechanisms, depending on their origin, oxidation status, composition, and additional xenobiotics. The exposure routes also seem to be a crucial criterion to investigate NPs’ toxicity. Further studies may give a greater consideration to environmentally relevant NPs, to get closer to field conditions undergone by aquatic organisms.

## Acknowledgements

We want to thank the LabEx COTE funding for supporting the PLASCOTE project (ANR 10-LABX-0045) and the University of Bordeaux for funding through its “inter-department” call 2021. We also thank the molecular biology platform of the EPOC laboratory for giving us access to the transcriptomic and enzymatic assays and PCR equipment, Amélie Vax-Weber for the search of possible THF traces in the NP-PS medium by GC-MS analysis, and Diana Kazaryan for obtaining TEM images of NP-L and NP-PS at the Bordeaux Imaging Centre electron microscopy facility (www.bic.u-bordeaux.fr/).

## Supporting Information

**Figure S1:**
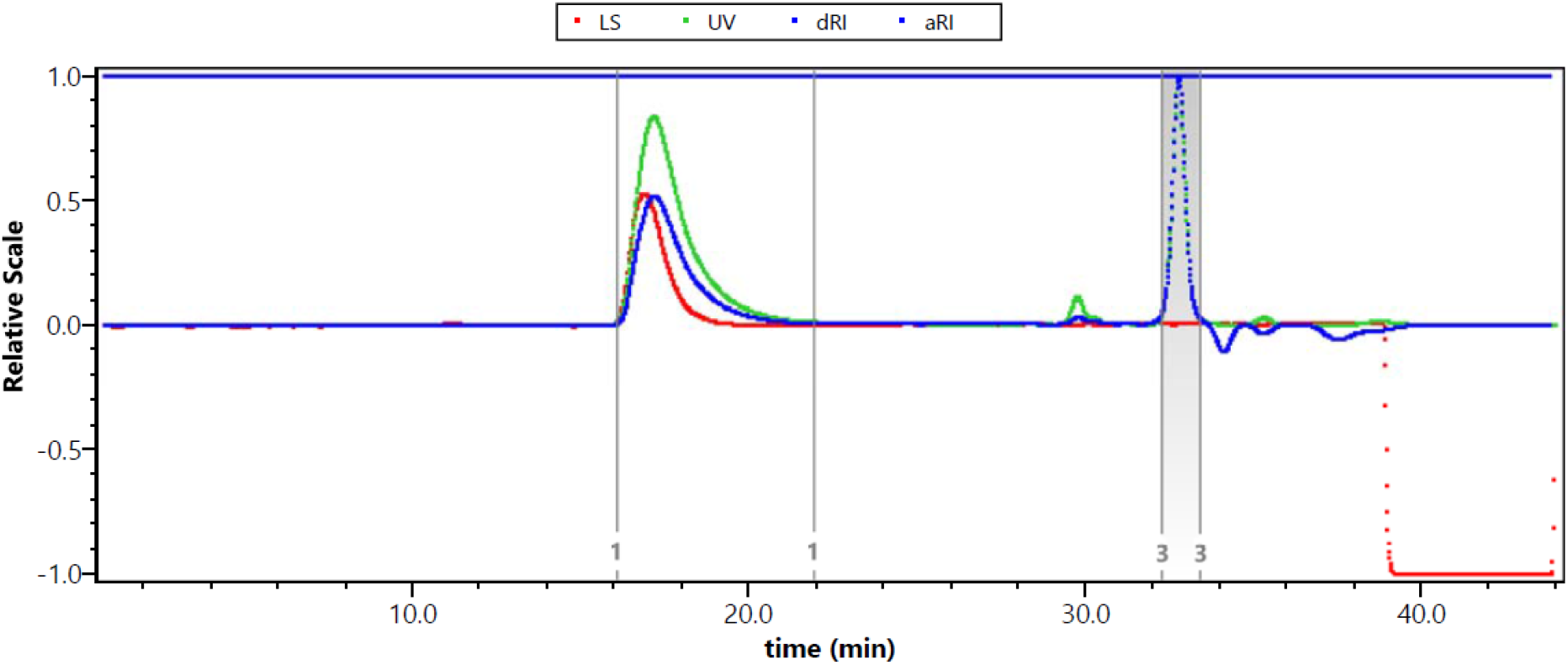
Size exclusion chromatogram of the commercial PS chains used to prepare NP model nanoplastics. The peak 1 on the three detectors (light scattering, UV and differential RI) is assigned to the sample yielding to *M*_n_=89 kg/mol, *M*_w_=196 kg/mol and *Ð*=*M*_w_/*M*_n_=2.21. Peak 3 is the flow marker.

**Figure S2:**
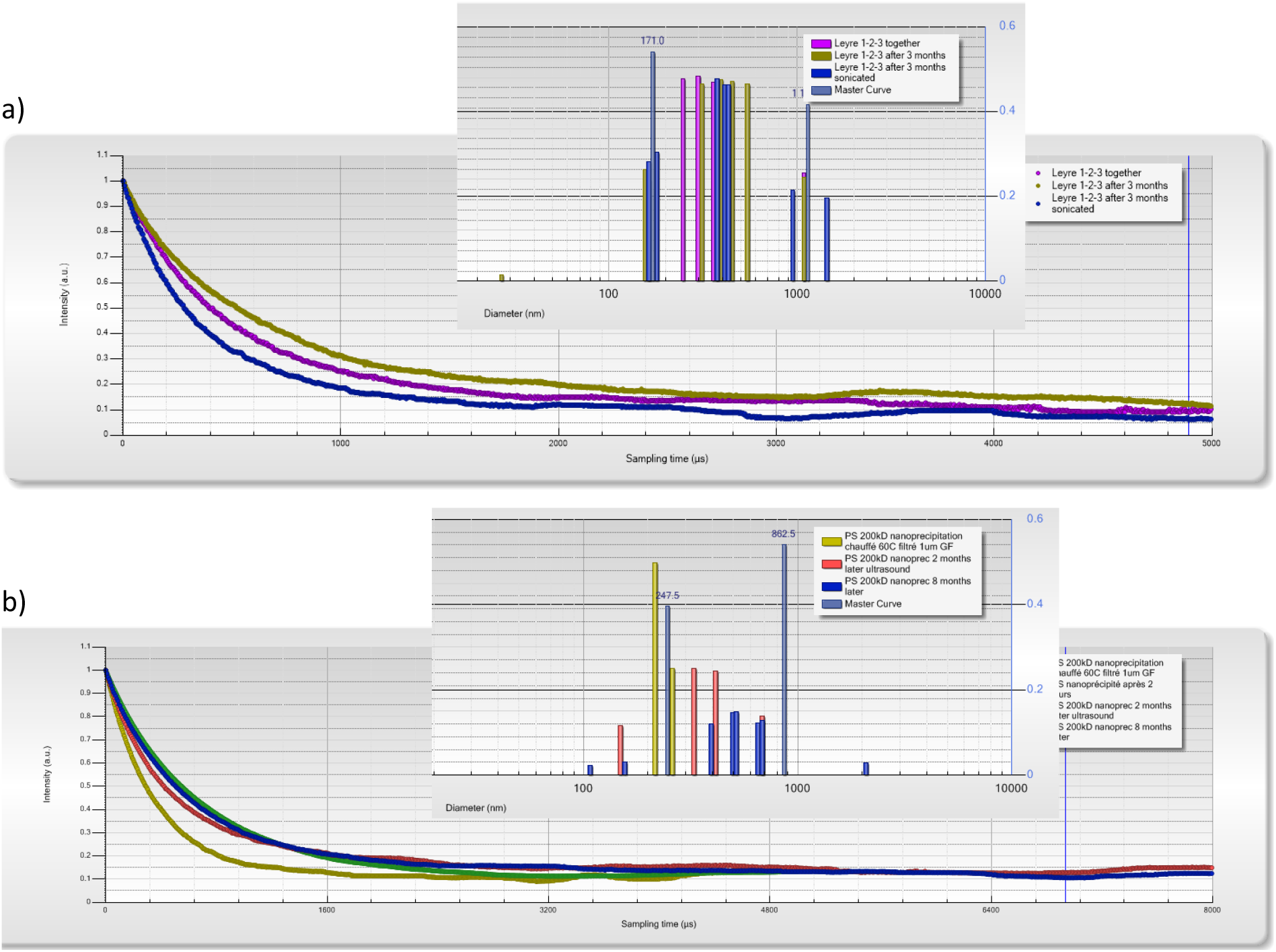
Kinetic study of colloidal stability by DLS (at 165° scattering angle, on a Vasco™ Flex remote-head backscattering DLS instrument, Cordouan Technologies, Pessac, France) of a) NP-L and b) NP-PS during several months after their preparation (when indicated, simple sonication with an ultrasound bath for 5 min was performed, before measurement). Insets show hydrodynamic size histograms obtained by the multimode Pade-Laplace fitting method.

**Figure S3:**
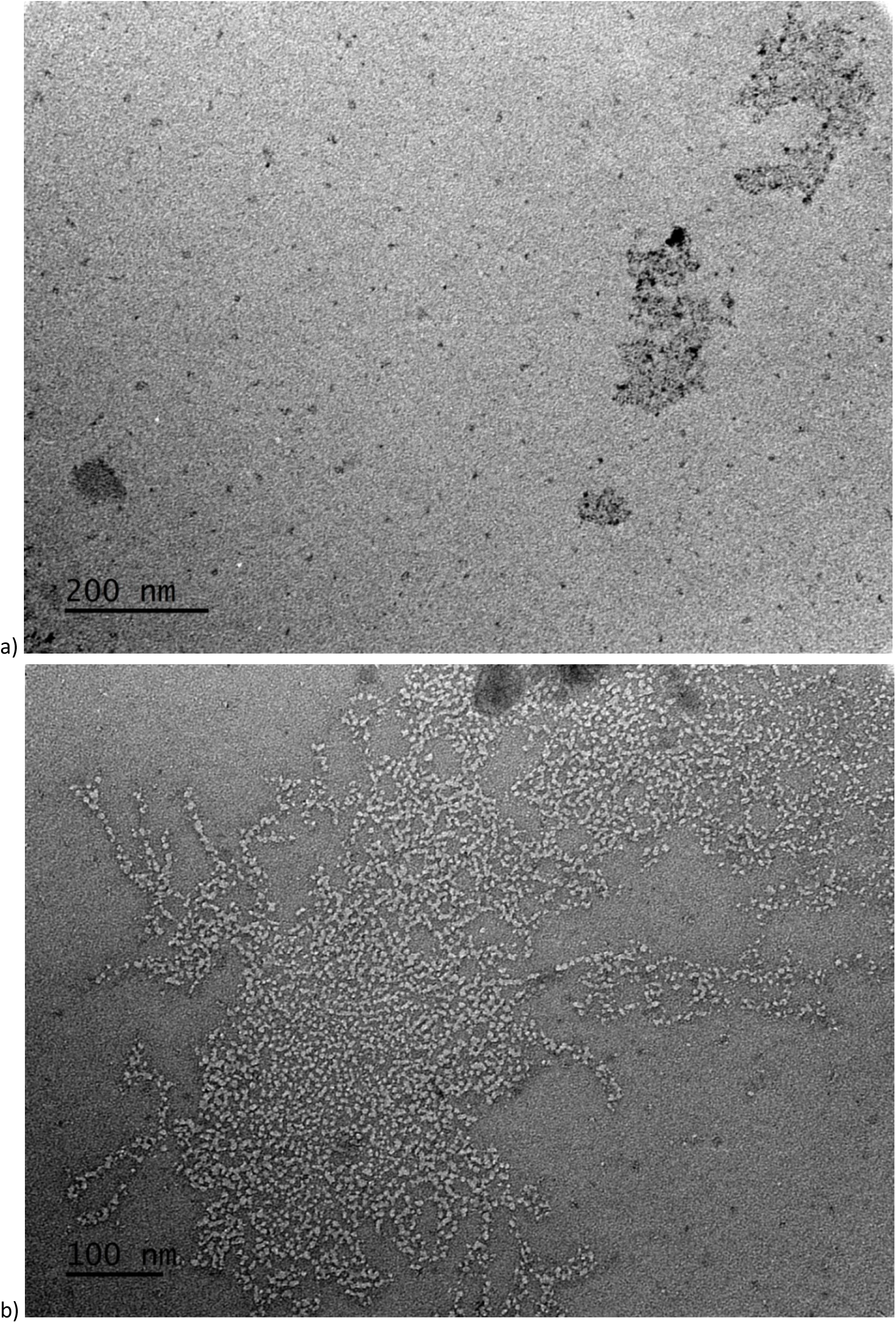
TEM image of a) NP-L and b) NP-PS (with samarium acetate staining specifically hydrophilic regions) showing nanoparticles of undefined morphologies in both cases.

**Figure S4:**
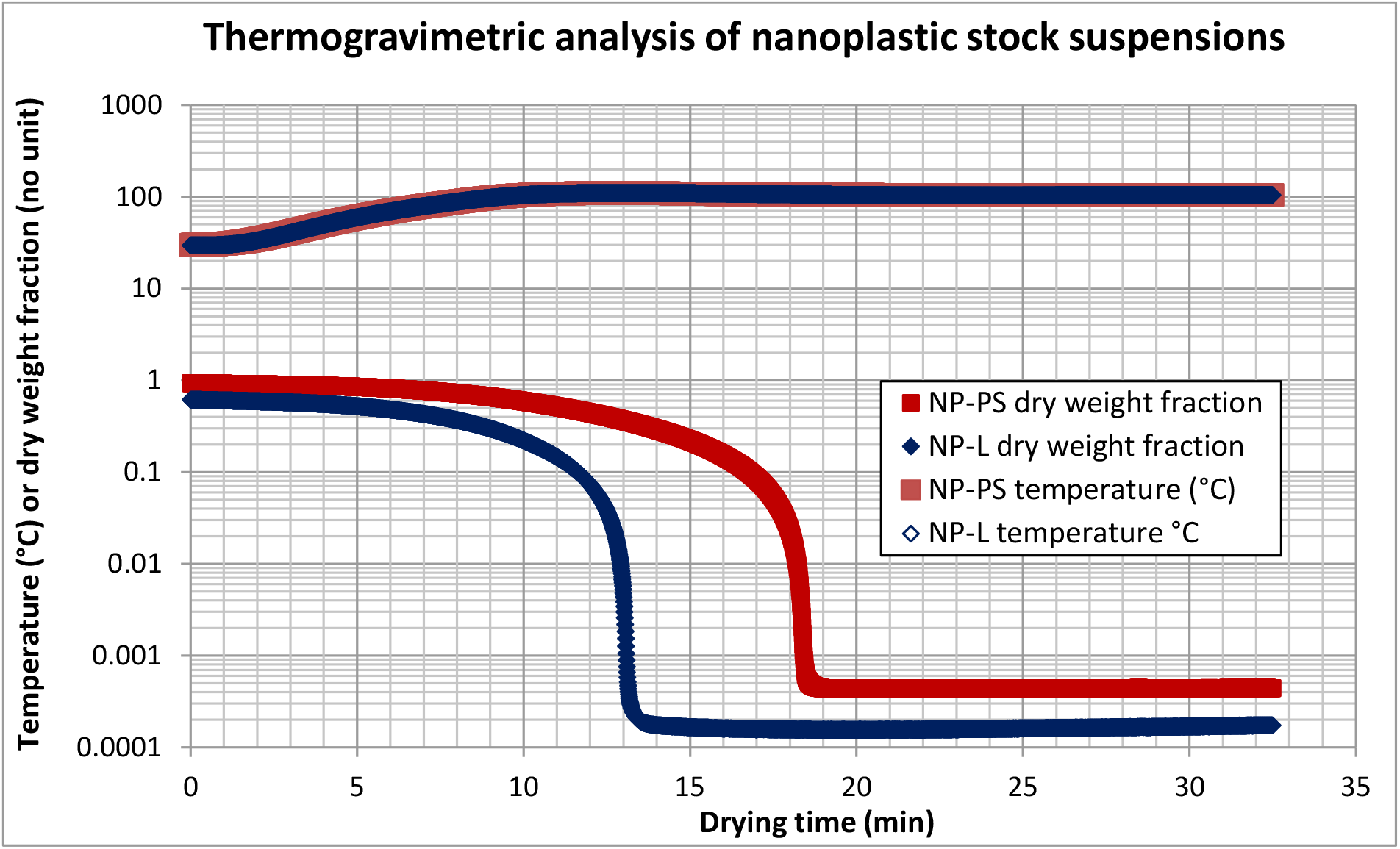
Thermogravimetry analyses (TGA) of a) NP-L and b) NP-PS suspensions yielding the weight concentrations of the stock suspensions: 147 mg/L for NP-L and 420 mg/L for NP-PS.

